# A bestiary of localized sequence rearrangements in human DNA

**DOI:** 10.1101/175943

**Authors:** Martin C. Frith, Sofia Khan

**Affiliations:** Artificial Intelligence Research Center, National Institute of Advanced Industrial Science and Technology (AIST), Tokyo, Japan; Graduate School of Frontier Sciences, University of Tokyo, Chiba, Japan; Computational Bio Big-Data Open Innovation Laboratory (CBBD-OIL), National Institute of Advanced Industrial Science and Technology (AIST), Tokyo, Japan

## Abstract

Genomes mutate and evolve in ways simple (substitution or deletion of bases) and complex (e.g. chromosome shattering). We do not fully understand what types of complex mutation occur, and we cannot routinely characterize arbitrarily-complex mutations in a high-throughput, genome-wide manner. Long-read DNA sequencing methods (e.g. PacBio, nanopore) are promising for this task, because one read may encompass a whole complex mutation. We describe an analysis pipeline to characterize arbitrarily-complex “local” mutations, i.e. intrachromosomal mutations encompassed by one DNA read. We apply it to nanopore and PacBio reads from one human cell line (NA12878), and survey sequence rearrangements, both real and artifactual. Almost all the real rearrangements belong to recurring patterns or motifs: the most common is tandem multiplication (e.g. heptuplication), but there are also complex patterns such as localized shattering, which resembles DNA damage by radiation. Gene conversions are identified, including one between hemoglobin gamma genes. This study demonstrates a way to find intricate rearrangements with any number of duplications, deletions, and repositionings. It demonstrates a probability-based method to resolve ambiguous rearrangements involving highly similar sequences, as occurs in gene conversion. We present a catalog of local rearrangements in one human cell line, and show which rearrangement patterns occur.

## Introduction

Genomes mutate and evolve by base substitutions, deletions, inversions, tandem duplications, and more complex events [1], up to large-scale chromosome shattering [2]. DNA sequence rearrangements contribute to a wide range of disorders, including developmental disorders (e.g. [3, 4]) and common diseases (e.g. [5]). In addition, somatic rearrangements serve as hallmarks in tumor genomes (e.g. [6, 7]).

The methods for studying structural variations have long been laborious, time-consuming or low resolution, e.g. fluorescence in situ hybridization (FISH), comparative genomic hybridization (CGH) or genome-wide SNP arrays (reviewed in [8]). Short-read DNA sequencing can provide high-throughput base-level resolution (e.g. [9, 10]); however, the read length limits the types of structural variation detected, and limits reliable alignment to repetitive regions [11], including subtelomeric regions that are known to harbor rearrangements [12].

A recent study surveyed “large” structural variation in 689 people, at *∼*5 kb resolution, by both long-insert and linked-read whole-genome sequencing [3]. On one hand, they found many novel “complex” variants: most (97%) are intrachromosomal, and they were classified into 16 types. On the other hand, their data cannot resolve micro-complexity (*<*5 kb), so they likely underestimate the complexity and diversity of complex variants.

Long-read DNA sequencing technologies provide read lengths to span tens of kilobases and beyond, which enables investigation of complex rearrangements spanned by one read. Long-read DNA sequencing has been used on its own [13, 14], or in combination with short reads [15], to investigate structural variation. The present tools for structural variant calling solely from long read data focus on insertions, deletions, and inversions (SMRT-SV [13]) and additionally translocations (MultiBreaks-SV [16] and PBHoney [17]).

In this study, we describe a method for finding arbitrarily-complex “local” re-arrangements, i.e. intrachromosomal rearrangements spanned by one DNA read. (We ignore pure deletions, which are relatively common and well-studied.) Our approach has two unusual features. First, we consider the ancestral state of each rearrangement, which is necessary in general to characterize them accurately. It also leads to a better understanding of “insertions”. Second, our approach is based on probabilities. There is often ambiguity in how to align rearranged fragments of a read, for example in gene conversion (replacement of a DNA segment by a duplicate of a similar segment). Such ambiguity can be resolved by calculating the probabilities of alternative alignments, based on the probabilities of substitutions and gaps due to sequencing error. These two features come together in an overlooked classical algorithm, the Repeated Matches Algorithm [18], which finds the most probable alignments of derived to ancestral sequences allowing for any number of duplications, deletions, repositionings, and insertions.

This method is applied to both MinION and PacBio DNA reads from the same human cell line. We describe error rates, and the troubling presence of dubious rearrangements that are likely artifacts of the DNA sequencing process. By semi-manual analysis, we extract 166 likely-real rearrangements, many of which are found in both MinION and PacBio. This reveals several recurring patterns of rearrangement.

As this manuscript was being finalized, a preprint appeared describing NGMLR and Sniffles, methods for finding structural variants in long reads (Sedlazeck et al. doi:10.1101/169557). Our study is largely complementary to theirs. Whereas Sniffles attempts to characterize variants automatically, we draw pictures of them and manually survey the types of rearrangement. We find intricate rearrangements, such as heptuplication, localized shattering, and gene conversion, not mentioned in NGMLR/Sniffles. Finally, NGMLR/Sniffles do not consider ancestral arrangements, and do not use fitted probabilities.

The remainder of this Introduction section describes the relevant aspects of DNA sequence evolution and probability-based alignment.

### Comparing rearranged sequences

In order to find rearrangements between two sequences, we must first determine “equivalent” positions between them, i.e. align them. To clarify this, let us consider how DNA sequences can mutate:

- Substitution of bases.
- Deletion of one or more consecutive bases.
- Duplication of sequence.
- Spontaneous generation: insertion of sequence not descended from any ancestor. This is unusual. An example is retrotranscription and insertion of RNA poly-A tails, which are not descended from any ancestral sequence.
- Repositioning of sequence segments, e.g. inversion or translocation.
- Insertion of sequence from another (e.g. viral) genome.

One mutational event may produce several of these effects, e.g. repositioning and deletion and duplication [1].

Let us now consider sequence evolution with complex mutations. Fig. 1 shows an ancestral sequence (**A**) evolving into two derived sequences (**B** and **C**). If we wish to find equivalent positions between **B** and **C**, this means identifying bases that descend from the same base in their most recent common ancestor. In general, this is hard: for example, it is hard to avoid an incorrect alignment between segment 3 in **B** and segment 4 in **C**.

**Figure 1.**
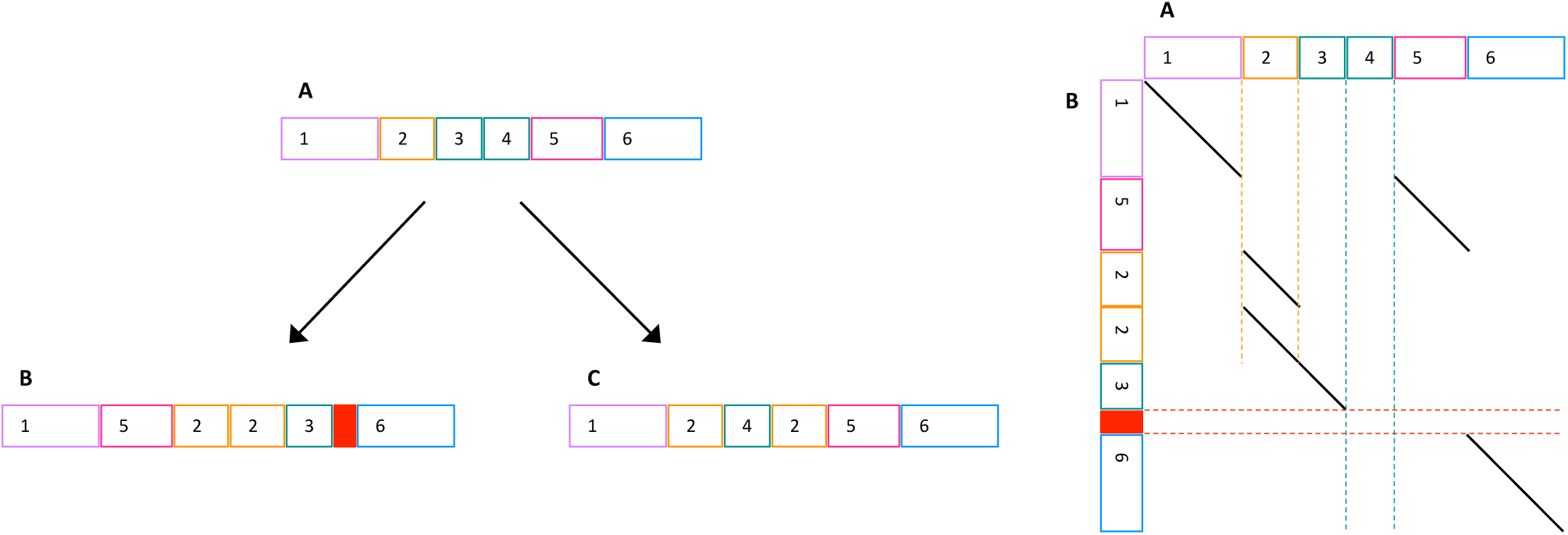
Left: sketch of sequence evolution by complex mutations. **A** is an ancestral sequence; **B** and **C** are derived. Same-color blocks (e.g. 3 and 4 in **A**) indicate similar sequences. The solid red block is “spontaneously generated”, i.e. not descended from any ancestral sequence. Right: alignment between **A** and **B**. Dashed lines indicate: duplication (vertical orange), deletion (vertical teal), and spontaneous generation (horizontal red).

The key observation is that it’s fundamentally easier to compare a derived sequence to an ancestral sequence, e.g. **B** to **A**. This is because *every* part of the derived sequence (apart from spontaneously-generated parts) is descended from a *unique* part of the ancestral sequence(s). Thus, a reasonable approach is to seek an optimal division of **B** into parts, where each part is aligned to a most-similar part of **A**. It may be objected that we rarely have an ancestral sequence, but this can be worked around using outgroups (see below).

### Classic probability-based alignment

The classic approach to sequence alignment, which ignores rearrangement, is based on probabilities [18]. First, we must determine the probabilities of each kind of substitution (e.g. `c`→ `t`), and insertion and deletion probabilities, for a specific kind of sequence comparison. Four examples are shown in Table 1. Then, we can align sequences *while considering these probabilities*: in other words, prefer alignments with higher probability [18]. It stands to reason that this approach will tend to produce more accurate alignments (than alignment by ad hoc criteria not based on the rates of substitution, insertion and deletion).

**Table 1.**
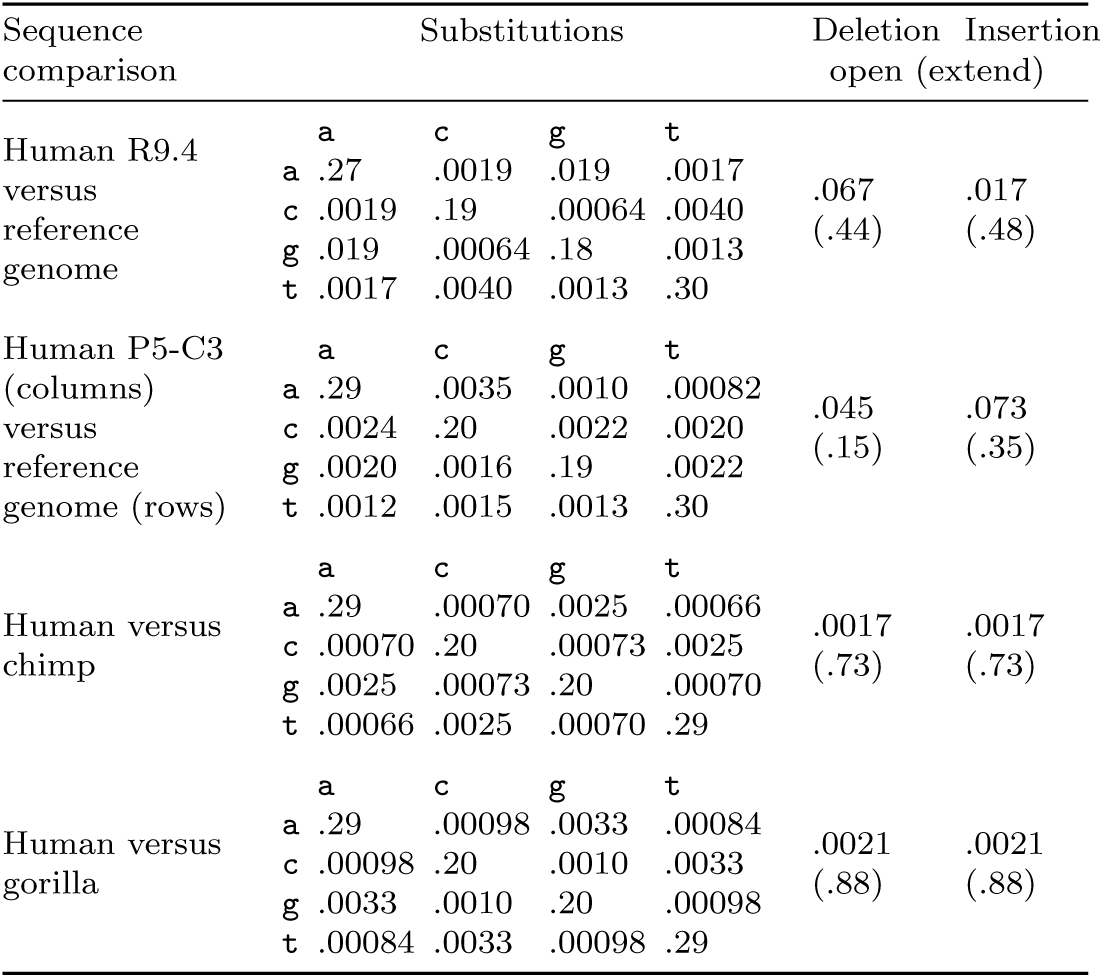
Probabilities of substitution, deletion, and insertion

### Probability-based alignment with rearrangements

Thus, we would like to find an optimal division of a derived sequence into parts, and optimal alignment of each part to ancestral sequence(s) based on probabilities of substitution, insertion and deletion. Fortunately, there is an algorithm to do exactly this: the Repeated Matches Algorithm (in section 2.3 of [18]). This algorithm seems to do more work than classic alignment (because it finds an optimal segmentation *and* alignment), but its computational cost is not significantly greater [18]. It is too slow for human-genome-scale data, so fast heuristic approximations are needed. One such heuristic is `last-split` [19].

### Ambiguity and alignment probabilities

Alignments have varying degrees of ambiguity. For example, suppose that our ancestral sequence has several identical copies of a retrotransposon, one of which has been retrotransposed (i.e. duplicated and inserted at a random location) in the derived sequence. While it is true that this insertion descends from a unique part of the ancestral sequence, this unique ancestral part is unknowable.

By using the Repeated Matches Algorithm’s probabilistic interpretation, we can quantify such ambiguity [19]. In the above retrotransposon example, the algorithm would align the insertion to an arbitrarily-chosen copy in the ancestral sequence, and annotate these bases with low correct-alignment probabilities (high “mismap” probabilities). This is useful, and the best that can be done.

## Materials and Methods

### DNA read data

The MinION (R9.4) dataset (14 million reads, 90 billion bases) is from `rel3` at https://github.com/nanopore-wgs-consortium/NA12878 (Jain et al. doi:10.1101/128835). The PacBio (P5-C3) dataset (30 million reads, 197 billion bases) is from accession SRX627421 [15]. Note that the PacBio reads are a few years older, so this may not be a fair comparison of the technologies as they are now.

### Alignment of DNA reads to the hg38 human genome

The two datasets, R9.4 and P5-C3, were analyzed separately in this and all following steps. First, the probabilities of substitution, insertion, and deletion were determined by `last-train`. Then, the reads were aligned to the genome by `last-split` (exact commands in the Supplement).

### Outgroup alignments

The `hg38` human genome was aligned to genomes of chimp [20] and gorilla [21]. The precise methods, and resulting alignments, are available at https://github.com/mcfrith/last-genome-alignments.

### Finding rearrangements

Rearrangements were found by these steps:

1. In each read separately, identify “local” rearrangements, i.e. intrachromosomal rearrangements encompassed by one read.
2. Cluster overlapping rearrangements (from multiple reads) into “rearranged regions”.
3. Discard each rearranged region that is not covered by an unbroken alignment of chimp or gorilla DNA to the reference human genome. This aims to get rearrangements where the reference human genome’s arrangement is ancestral. (It could fail to do so, if the derived arrangement was present in the common ancestor of these species.)

The software is available at: https://github.com/mcfrith/local-rearrangements.

### Re-aligning non-orphan rearrangements

The rearrangements include many orphans present in just one read (more likely to be artifacts), and fewer non-orphans present in multiple reads (more likely to be real). In order to characterize the non-orphans more accurately, these reads were re-aligned to the genome more slowly-and-sensitively, by using `lastal` options `-m50 -d90` (exact commands in the Supplement).

### Genome annotations

Annotations of the `hg38` human genome were taken from its `refFlat.txt` [22], `rmsk.txt` [23], and `simpleRepeat.txt` [24] files, in the UCSC genome database [25].

## Results

### Sequencing error rates

The rates of substitution, insertion, and deletion relative to a reference human genome (`hg38`) were estimated with `last-train` [26]. These rates include both sequencing errors and real genome differences, but greatly exceed the well-established rates of real difference, so are mostly error (Table 1).

Both R9.4 and P5-C3 have more indel than substitution errors. R9.4 has more deletions (3.9×) than insertions. P5-C3 has more insertions (4.3×) than R9.4, but they tend to be shorter. It has fewer deletions (0.7×), also shorter.

A preliminary result for R9.4 (not shown) had an almost-symmetric substitution matrix, therefore the training was redone assuming a symmetric matrix (Table 1). P5-C3 lacks this symmetry. R9.4 has higher rates of transition errors, especially `a↔g`.

We also estimated the rates of substitution, deletion and insertion for chimp and gorilla DNA versus human (in order to make outgroup alignments). Compared to the long-read sequencing errors, there are much fewer insertions and deletions, but they tend to be longer. The rates of transition (`a↔g` and `c↔t`) are higher than those of P5-C3, but lower than those of R9.4. The transversion rates are mostly lower than those of the long reads.

### Orphan rearrangements

Because each dataset has many-fold coverage of the genome, we expect each rearrangement to be covered by multiple DNA reads. Surprisingly, however, there are many orphan rearrangements appearing in just one read.

The R9.4 dataset has 33 695 rearranged regions, of which 31 600 have just one rearranged read. Tandem duplications (TDs) are frequent: 29 339 of the orphans, and 1169 of the non-orphans, are TDs. A (pseudo)random sample of TD and non-TD orphans is shown in Fig. 2. The non-TD orphans often have a zigzag pattern, implying triplication.

**Figure 2.**
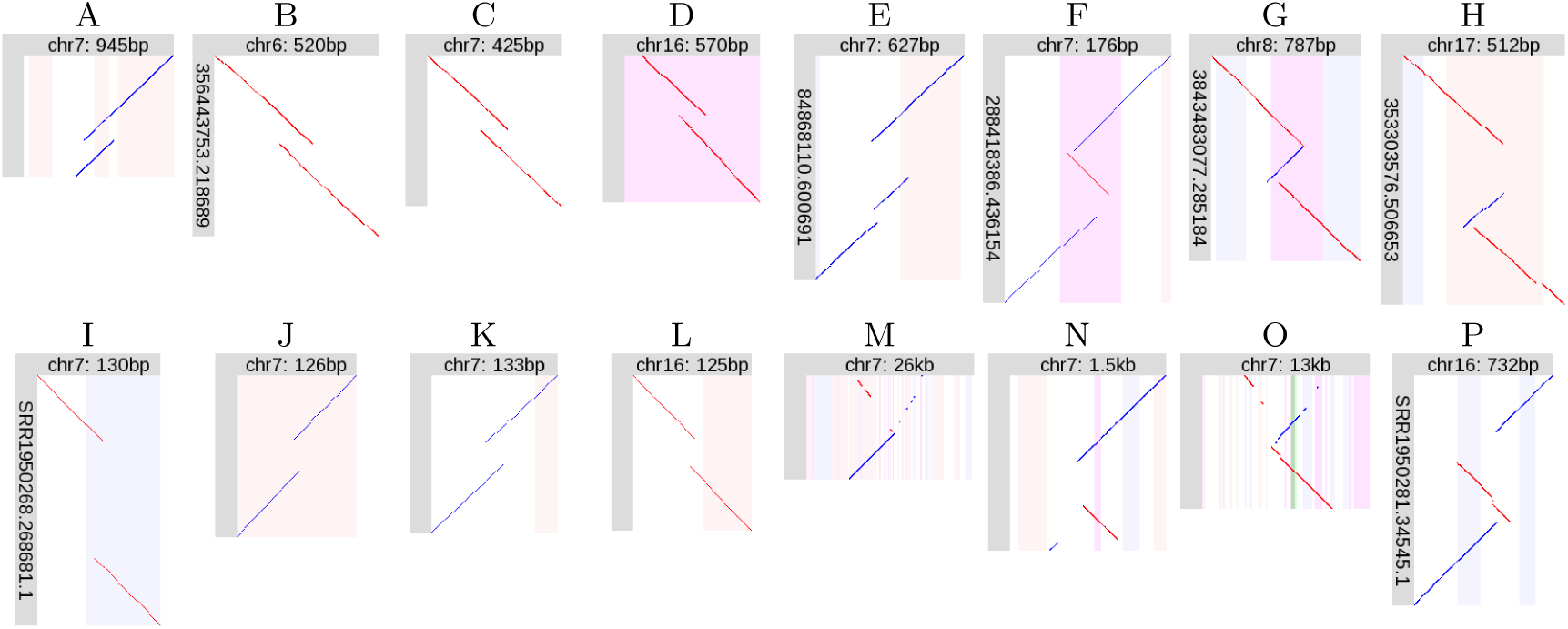
Orphan rearrangements. **A**–**D**: R9.4 tandem duplications (TDs). **E**–**H**: R9.4 non-TD rearrangements. **I**–**L**: P5-C3 tandem duplications. **M**–**P**: P5-C3 non-TD rearrangements. Diagonal lines indicate alignments between a segment of a DNA read (vertical), and a segment of the reference human genome (horizontal). Red lines indicate same-strand alignments; blue lines indicate opposite-strand alignments. The vertical stripes indicate features in the reference genome; pink: forward-strand trans-posable element, blue: reverse-strand transposable element, purple: low-complexity or tandem repeat, green: exon.

The P5-C3 dataset has 196 527 rearranged regions, of which 183 653 have just one rearranged read. TDs are again frequent: 154 487 orphans (and 3143 non-orphans) are TDs. A sample of these orphans is also shown in Fig. 2. Each TD orphan has a very small duplication (compared to R9.4), flanking an unaligned part of the read. The non-TD orphans often have a bizarre fountain-like appearance (Fig. 2 M,O), with interspersed fragments from opposite strands.

A plausible explanation of these orphan rearrangements is that they are artifacts of the DNA sequencing process.

### Supported rearrangements

Non-orphan rearrangements were analyzed further. Most non-orphans are tandem duplications and/or inside tandem repeats: the latter is not surprising, because tandem repeats have high rates of tandem duplication and gene conversion [27]. To avoid manual inspection of numerous similar rearrangements, we discarded tandem duplications (but not triplications etc.), and rearranged regions *>* 50% covered by tandem repeats.

For the R9.4 dataset, this left 164 rearranged regions, which were manually inspected (Supplemental figures). A few were judged unclear or dubious (Table 2), usually due to doubt that the reference human genome’s arrangement is ancestral. (Gaps ≤ 50 bp were allowed in the ape-human alignments, and sometimes these gaps coincided suspiciously with small rearrangements.)

**Table 2.** Number of rearrangements of each type

| Type | Total | Found with |  |
| --- | --- | --- | --- |
|  |  | R9.4 | P5-C3 |
| Non-tandem duplication (same strand) | 26 | 26 | 14 |
| Non-tandem duplication (opposite strand) | 21 | 21 | 13 |
| Non-tandem double duplication | 2 | 2 | 0 |
| Inverted tandem duplication | 6 | 6 | 5 |
| Inversion | 8 | 8 | 3 |
| Inversion next to deletion | 12 | 12 | 6 |
| Inversion in deletion | 8 | 8 | 1 |
| Non-tandem duplication (same strand) with deletion | 6 | 6 | 1 |
| Non-tandem duplication (opposite strand) with deletion | 6 | 6 | 2 |
| Localized shattering | 4 | 4 | 1 |
| Conversion | 4 | 4 | 1 |
| Tandem multiplication | 54 | 42 | 24 |
| Unique | 9 | 8 | 2 |
| TOTAL | 166 | 153 | 73 |
| Unclear | 21 | 2 | 19 |
| Dubious | 36 | 9 | 29 |

For the P5-C3 dataset, it is harder to separate plausible from dubious rearrangements. There are more dubious rearrangements, and the fountain-like ones are especially troublesome, because they are large (e.g. 10 kb) and often overlap other rearrangements. We therefore applied a stricter criterion, retaining only rearranged regions that have at least four reads with non-TD rearrangements. This left 121 rearranged regions (Supplemental figures), of which 48 were judged unclear or dubious (Table 2), usually because they have multiple overlapping fountain-like rearrangements.

Ultimately, this semi-manual analysis produced 166 local rearrangements judged to be clear and reliable (Table 2). One argument for their reliability is that each case has more than one read with the *same* rearrangement (Supplemental figures). An example is shown in Fig. 3A: this shows two reads, one from each DNA strand, with the same tandem heptuplication. In contrast, there are many cases where fountain-like rearrangements overlap each other, but they are different (e.g. Fig. S250, Fig. S274).

**Figure 3.**
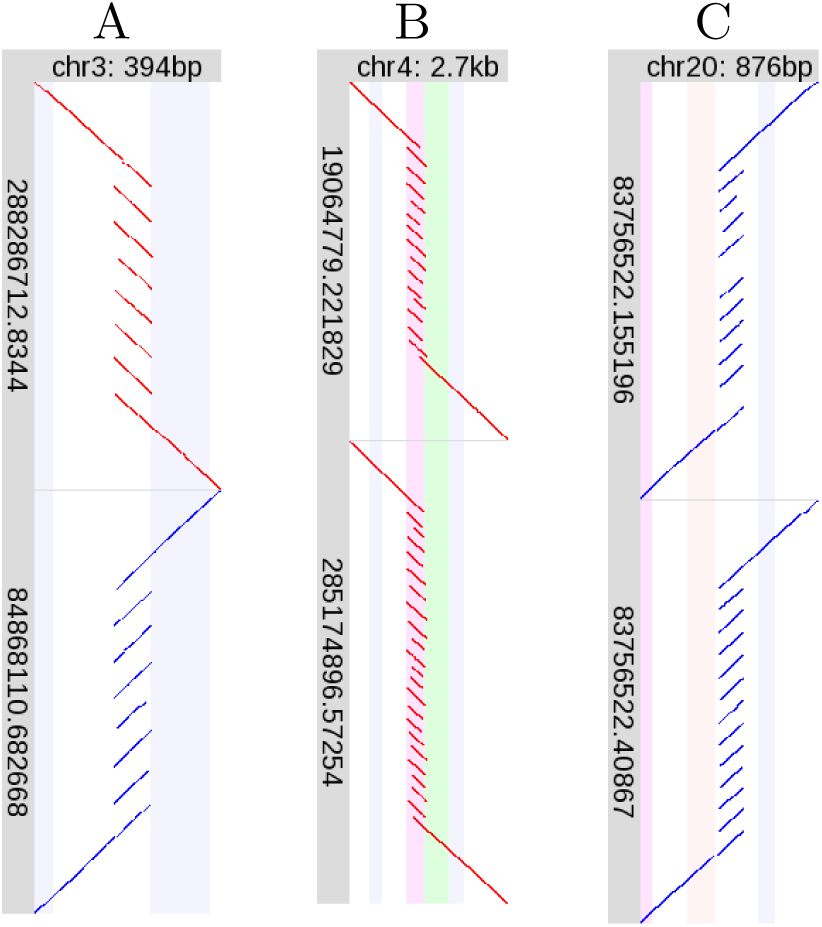
Examples of tandem multiplication. Please see the description of Fig. 2. Here, each of **A**, **B** and **C** shows two R9.4 DNA reads, one above the other.

Another argument for the reality of these rearrangements is that 60 of them are found by both R9.4 and P5-C3, independent sequencing technologies. In fact, the P5-C3 rearrangements are mostly a subset of the R9.4 rearrangements, suggesting that with R9.4 we find almost all local rearrangements present in this genome. The main exception is the tandem multiplication category, where P5-C3 has 12 rearrangements not found in R9.4 (Table 2). This is related to our removal of rearrangements in tandem repeats: most of the 12 would be in tandem repeats, except for one or two reads with dubious alignments that enlarge the rearranged region (e.g. Fig. S184). On the other hand, the reason for finding only a subset of the rearrangements in P5-C3 is (at least partly) the stricter criterion used to discard dubious rearrangements.

### Zygosity

A further argument for the reality of these rearrangements is from zygosity. Since the DNA reads are from diploid cells (with two copies of each chromosome), if a rearrangement is real, it should be either homozygous (in both copies of the chromosome), or heterozygous (in just one copy). By examining all the DNA reads that align to a rearranged region, we can judge its zygosity.

Many of the rearranged regions have a clear zygosity. For example, the tandem multiplication in Fig. 3A is heterozygous: about half the R9.4 DNA reads align straight across this region without rearrangement (Fig. S165). This is confirmed by P5-C3 reads (Fig. S166). In contrast, Fig. 3B shows two different rearrangements in the same region, implying that both chromosomes are rearranged, so there should be no unrearranged DNA reads here. Indeed, there are none (Fig. S169). Some homozygous examples are: Fig. 4H (Fig. S93), Fig. 4L (Fig. S106), Fig. 4N (Fig. S121), and Fig. 4P (Fig. S141). Some heterozygous examples are: Fig. 3C (Fig. S187), Fig. 4A (Fig. S2), Fig. 4B (Fig. S8), and Fig. 4J (Fig. S117). In some cases the zygosity is less clear, due to imperfect and ambiguous alignments.

**Figure 4.**
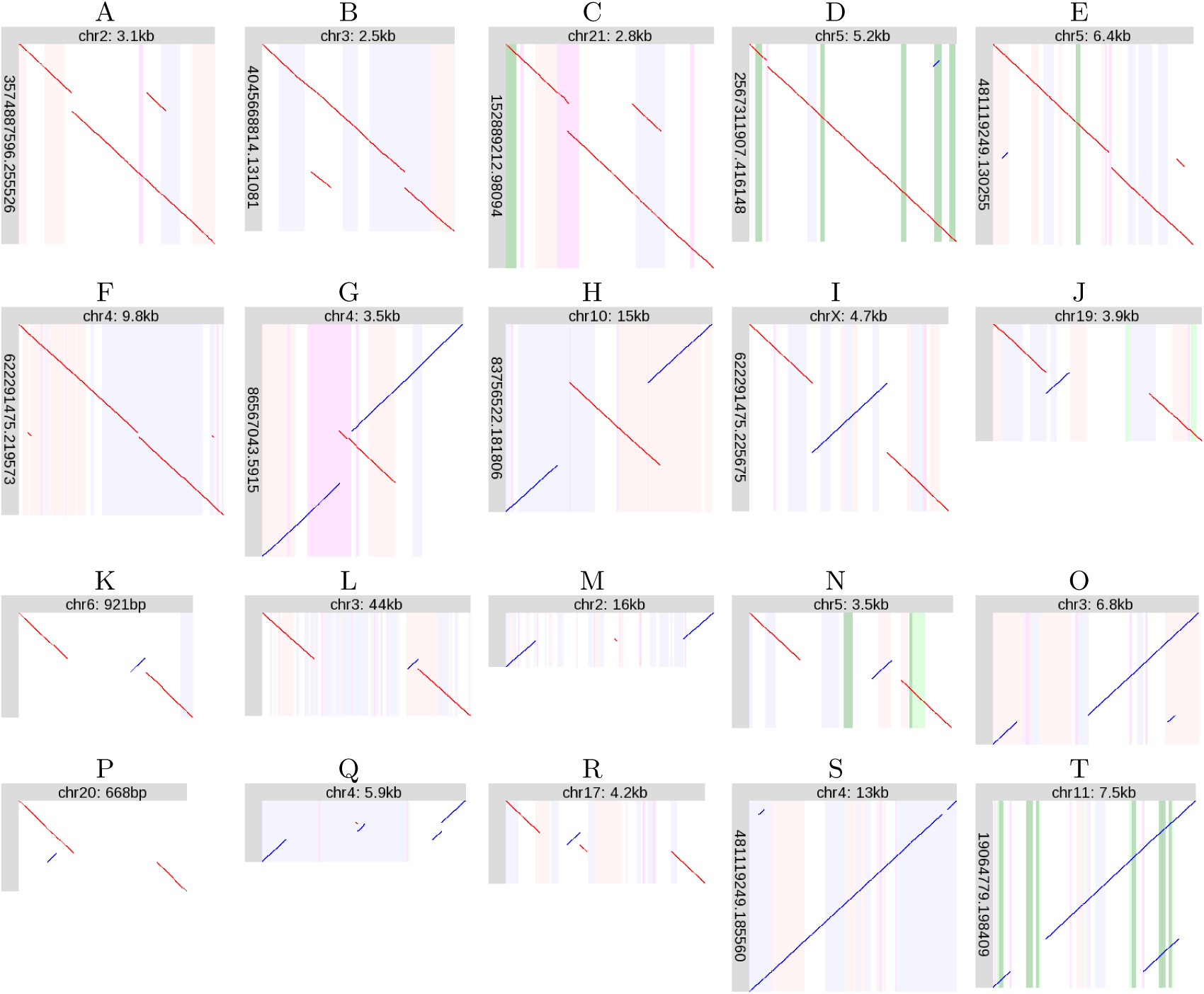
Examples of recurring rearrangement patterns. Diagonal lines indicate alignments between a segment of a DNA read (vertical), and a segment of the reference human genome (horizontal). Red lines indicate same-strand alignments; blue lines indicate opposite-strand alignments. The vertical stripes indicate features in the reference genome; pink: forward-strand transposable element, blue: reverse-strand transposable element, purple: low-complexity or tandem repeat, green: exon, dark green: protein-coding sequence. Some of the alignments (diagonal lines) are tiny: it may help to view this on a screen and zoom in.

### Recurring patterns of rearrangement

Many of the rearrangements strikingly resemble each other, i.e., there are recurring rearrangement motifs. Some of these motifs (generally, the simpler ones) match complex-variant classes from the limited-resolution study of Collins et al. [3].

#### Tandem multiplication

The most frequent type of rearrangement is tandem multiplication (Table 2). The degree of multiplication varies greatly, e.g. duplication (most common), heptuplication (Fig. 3A), and more (Fig. 3 B–C). The degree of multiplication is sometimes unclear, especially in tandem repeats: this is presumably due to ambiguity in alignment of tandemly-repeating sequences. The degree is often heterozygous, e.g. Fig. 3B shows a smaller allele with *∼*14-fold multiplication and a larger allele with *∼*22-fold multiplication.

The tandem multiplication in Fig. 3B is immediately upstream of the transcription start site of LINC01596 (long intergenic non-protein coding RNA 1596), so is likely to have a profound effect on the expression of this transcript.

Fig. 3C illustrates a problem that occurs with all types of rearrangement, but especially tandem multiplications. While the lower DNA read shows 12-fold tandem multiplication, the upper read has two missing fragments: these two parts of the read are unaligned. A plausible explanation is that the alignment method simply failed to align these fragments, perhaps because of sequencing errors. Such missing fragments can often be rescued by re-aligning the DNA read with more slow-and-sensitive heuristics.

Most of the tandem multiplications are near telomeres (Fig. S1), regions known to be rearrangement hot spots [12].

#### Non-tandem duplication

Another frequent pattern is non-tandem, but localized, duplication (Table 2). The duplicated segment can be inserted in its original orientation (Fig. 4 A–C), or in the opposite orientation (Fig. 4D). These rearrangement patterns were reported previously [15].

There are two cases of double non-tandem duplication (Fig. 4 E–F). In both cases, two non-adjacent segments have been duplicated, and inserted adjacent to each other roughly half-way between the duplication sources.

### Inverted tandem duplication

Another recurring pattern is inverted tandem duplication (Fig. 4G). This superficially resembles the zigzag pattern of orphan rearrangements (Fig. 2 F–H), but is duplication rather than triplication.

There are some near-tandem inverted duplications (e.g. Fig. S63), so perhaps there is no sharp distinction between non-tandem opposite-orientation duplications (Fig. 4D) and the present category.

#### Inversion

Inversions can occur without deletion (Fig. 4I), but more often occur next to (Fig. 4 J–L) or in/between (Fig. 4 M–N) deletions. This fits with previous reports that inversions are often flanked by gaps [28, 29, 30].

The example in Fig. 4N deletes the second-last protein-coding exon of SPINK14 (serine peptidase inhibitor, Kazal type 14) [15]. Examination of all reads aligned here (Fig. S121) indicates that this rearrangement is homozygous.

The inversion in Fig. 4H is flanked by oppositely-oriented L1PA7 LINEs, suggesting that it was caused by homologous recombination between these LINEs. Fig. 4H shows a small deletion to the left of the inversion, and a small duplication to the right: this is gene conversion (replacement of a sequence by a duplicate of a similar sequence), which is a typical effect of homologous recombination.

#### Non-tandem duplication with deletion

Another recurring pattern is non-tandem (but localized) duplication, combined with deletion at the duplication target site. Here too, the duplicated segment can be inserted in its original orientation (Fig. 4O), or in the opposite orientation (Fig. 4P).

#### Localized shattering

A more complex rearrangement motif is “localized shattering” (Fig. 4 Q–R). This is so named because it looks like the ancestral sequence was shattered into multiple fragments, of which some were lost, and the others rejoined in random order and orientation. Accordingly, this motif exhibits deletion and repositioning but not duplication. Another diagnostic feature is that some of the rearranged fragments come from adjacent parts of the ancestral sequence. If the fragments came from randomly and independently chosen parts of the ancestral sequence, it is unlikely that they would be adjacent. Shattering naturally explains this adjacency.

All of these diagnostic features are also present in the inversion-next-to-deletion pattern, which might thus be regarded as a special case of localized shattering.

#### Conversion

Another well-known pattern of DNA sequence change is gene conversion, where one sequence tract is replaced by a duplicate of a similar tract (Fig. 4 S–T).

The example in Fig. 4T involves protein-coding genes. The three green stripes on the left indicate the three exons of HBG1 (hemoglobin subunit gamma 1), and those on the right indicate HBG2 (hemoglobin subunit gamma 2). Here, the first two exons of HBG1 have been replaced by the equivalent exons of HBG2. Because HBG1 and HBG2 are very similar, this changes just one amino acid: a threonine is converted to an isoleucine. This substitution is present in dbSNP (rs1061234).

At the DNA level, the segments of HBG1 and HBG2 that participate in this conversion differ by just one length-4 indel and a dozen substitutions. (The exact number of substitutions depends on the exact endpoints of the conversion, which are highly ambiguous.) Thus, these DNA reads could all-too-easily be aligned to the HBG locus co-linearly, without rearrangement. This example showcases the power of probability-based alignment, which calculates that the rearranged alignment is much more probable. The most informative alignment part (of one read) is shown in Fig. 5: a co-linear alignment would align this part of the read to HBG1.

**Figure 5.**
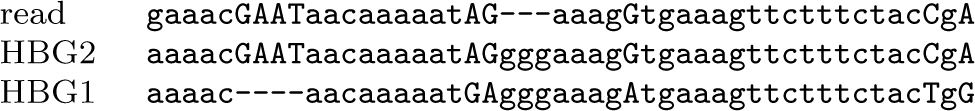
A segment of a DNA read aligned to HBG1 and HBG2. It is likely that this part of the read is paralogous to HBG1, and should rather be aligned to HBG2.

### Unique rearrangements

Some rearrangements were judged to be unique, and not follow any recurring pattern. The eight cases found in the R9.4 dataset are described here.

#### Inverted tandem duplication with deletion

Fig. 6A shows an inverted tandem duplication with adjacent deletion. The duplication and deletion have roughly the same size, suggesting gene conversion. However, the duplicated and deleted sequences lack similarity, which argues against conversion.

**Figure 6.**
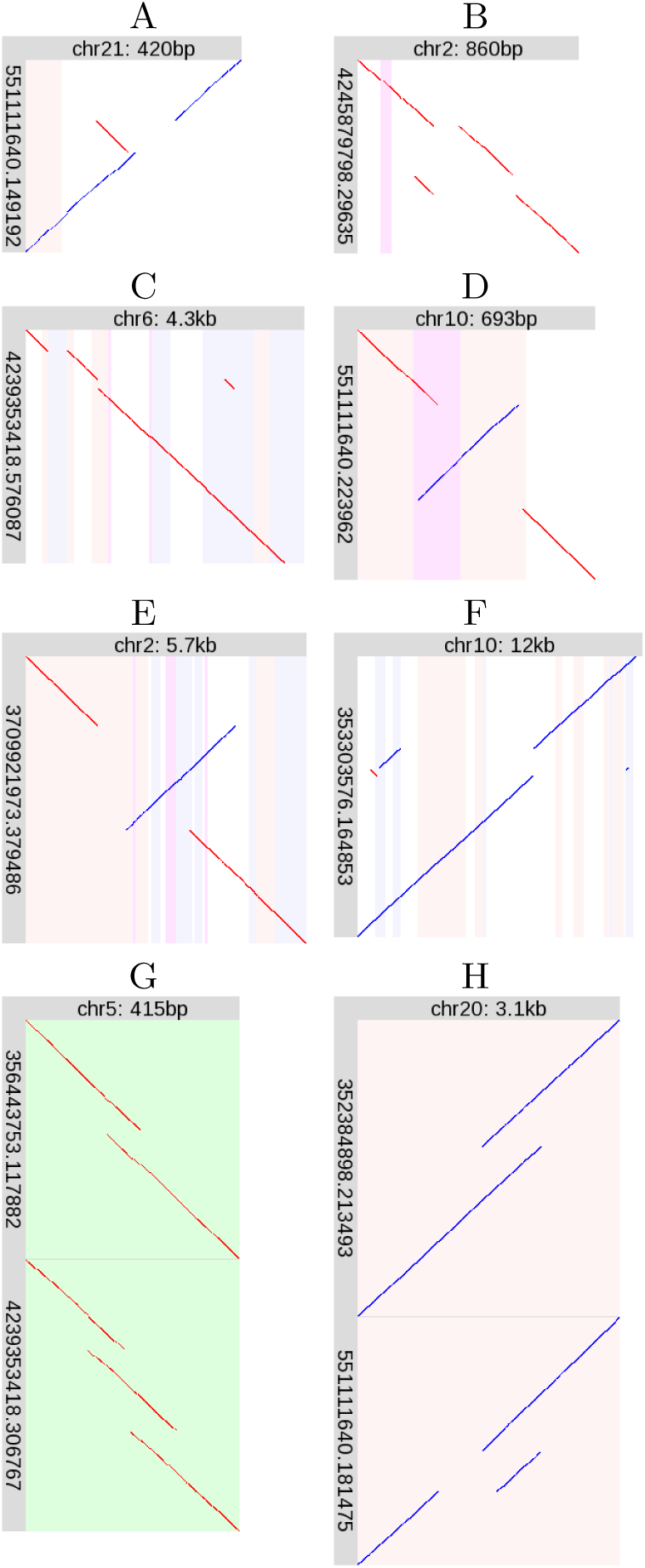
Unique rearrangements. Please see the description of Fig. 4.

#### Non-tandem duplication next to deletion

Fig. 6B shows a non-tandem duplication, combined with a deletion adjacent to the duplication *source*.

#### Non-tandem duplication with Alu deletion

Fig. 6C shows a non-tandem duplication, and near the duplication target site is a precise (or near-precise) deletion of an AluY element. Perhaps this deletion occured separately and independently of the duplication. Precise deletion of transposable elements is unusual [31]: maybe the reference human genome’s condition is not ancestral, even though it is shared by chimp and gorilla (Fig. S157).

#### Inversion with tandem repeat expansion

Fig. 6D shows an inversion that overlaps a period-45 tandem repeat (purple stripe). The tandemly repeated region has been partly duplicated. Since tandem repeats frequently expand, perhaps this inversion and expansion occurred independently.

#### Inversion with deletion and duplication

Fig. 6E superficially resembles inversion and gene conversion by homologous recombination (Fig. 4H). However, the duplicated and deleted regions are dissimilar and of different lengths (the deletion is *∼*600 bp and the duplicated region is *∼* 930 bp), which argues against conversion and recombination. Therefore, we suggest this rearrangement should be classified separately. Though it is unique in our study, this delINVdup pattern has been seen in other human DNA [32].

#### Multiple non-tandem duplication with conversion

Fig. 6F shows non-tandem duplication, with three duplicated fragments. The genomic sources of the two left-hand fragments are separated by 116 bp, and the right-hand fragment’s source is 114 bp. Moreover, the left-hand separation and right-hand fragment are in equivalent regions of Alu elements, suggesting that conversion contributed to this complex rearrangement.

#### Heterozygous adjacent tandem duplications

Fig. 6G shows two (heterozygous) rearrangements (relative to the reference genome) at the same location, in the 3’UTR of AHRR (aryl-hydrocarbon receptor repressor). One is a tandem duplication, the other is two adjacent tandem duplications: the latter pattern is unique in this study.

#### Tandem duplication next to heterozygous deletion

Fig. 6H shows two (heterozygous) rearrangements at the same location, within centromeric alpha satellite DNA. It appears that both rearrangements share the same tandem duplication, but one has an adjacent deletion.

### Repeat associations

Many of the rearrangements are strikingly associated with repeats. Firstly, tandem multiplications often overlap tandem repeats (e.g. Fig. 3B), as expected.

Secondly, some inversions and conversions are attributable to homologous recombination between repeats, also as expected. Fig. 4S shows conversion between L1PA3 LINEs. In Fig. 4J, the inverted segment is near-precisely flanked by oppositely-oriented AluSx SINEs, though this does not explain the deletion.

Thirdly, the target sites of some non-tandem duplications coincide with simple repeats (Fig. 4 C–E). An explanation may lie in the fact that simple repeats can be fragile [27].

Fourthly, rearrangement edges often coincide with edges of transposon fragments. In Fig. 4G, the duplication’s right end coincides with the 3’-end of an L1 LINE fragment (that lacks the 3’-most 5729 bases of a full-length L1). In Fig. 4N, the rightmost deletion edge is *∼* 6 bp downstream of the 5’-end of a (5’-truncated) MIR. The inverted fragment’s right edge is *∼*20 bp downstream of the 3’-end of a (3’-truncated) MamSINE1. In Fig. 4L, the deletion’s left edge is *∼*4 bp upstream of the 3’-end of a MIR (that lacks the 3’-most 126 bp of a full-length MIR). In Fig. 4M, the leftmost deletion edge is *∼*3 bp upstream of the 5’-end of an L1 (that lacks the first 6216 bp of a full-length L1). The rightmost edge is *∼*30 bp upstream of the 5’-end of an L2 (that lacks the first 3303 bp of a full-length L2). There are similar coincidences in Fig. 4 E,O,R and Fig. 6D.

Some tandem multiplications also coincide with edges of transposon fragments. The multiplication in Fig. 3A is next to the 3’-end of a (3’-truncated) MLT1H2 LTR. The multiplication in Fig. 3C sits neatly between a MER103C transposon (left) and an L2 LINE (right).

The duplication source in Fig. 4C near-perfectly overlaps an MLT1B LTR, suggesting retrotransposition of this element. This LTR has many mutations relative to the MLT1B consensus sequence, which enables unambiguous alignment, but suggests that it has lost retrotranspositional activity.

### Retrogene reduplication

The non-tandem duplication in Fig. 4B perhaps warrants further description. The duplication source is in a white region flanked by blue stripes. This white region is a retrotransposed copy of RPL23A (ribosomal protein L23a) [33], which was inserted in a Charlie5 transposon (blue stripes). Thus, the rearrangement in Fig. 4B is a reduplication of most of this retrogene.

There are many retrotransposed RPL23A copies in the genome [33], so we may wonder if this alignment is correct and unambiguous. This retrogene (the white region in Fig. 4B) has 90.2% identity to its parent mRNA [33]. The inserted part of the DNA read in Fig. 4B aligns here with 192 matches, 15 mismatches, and 64 gaps. This alignment has much higher probability than the second-best hit (187 matches, 20 mismatches, 82 gaps), to a retro-RPL23A on chromosome 19 with 97.7% identity to the parent mRNA. The insertion aligns to the parent mRNA with 186 matches, 21 mismatches, and 85 gaps, suggesting that it is not a direct retrotransposition of the parent gene.

### Rearrangement lengths

The rearrangement lengths (span in ancestral genome) vary between 50 and 20 000 bases (Fig. 7). The upper bound is limited by the read lengths. The lower bound presumably reflects the difficulty of aligning short, error-prone sequence fragments: shorter rearrangements exist [30]. Since the figure shows *ancestral* lengths, it is not surprising that tandem multiplications tend to be shorter, and rearrangements involving deletion tend to be longer. The conversions are longer-range than most of the other rearrangements (but there are just four of them).

**Figure 7.**
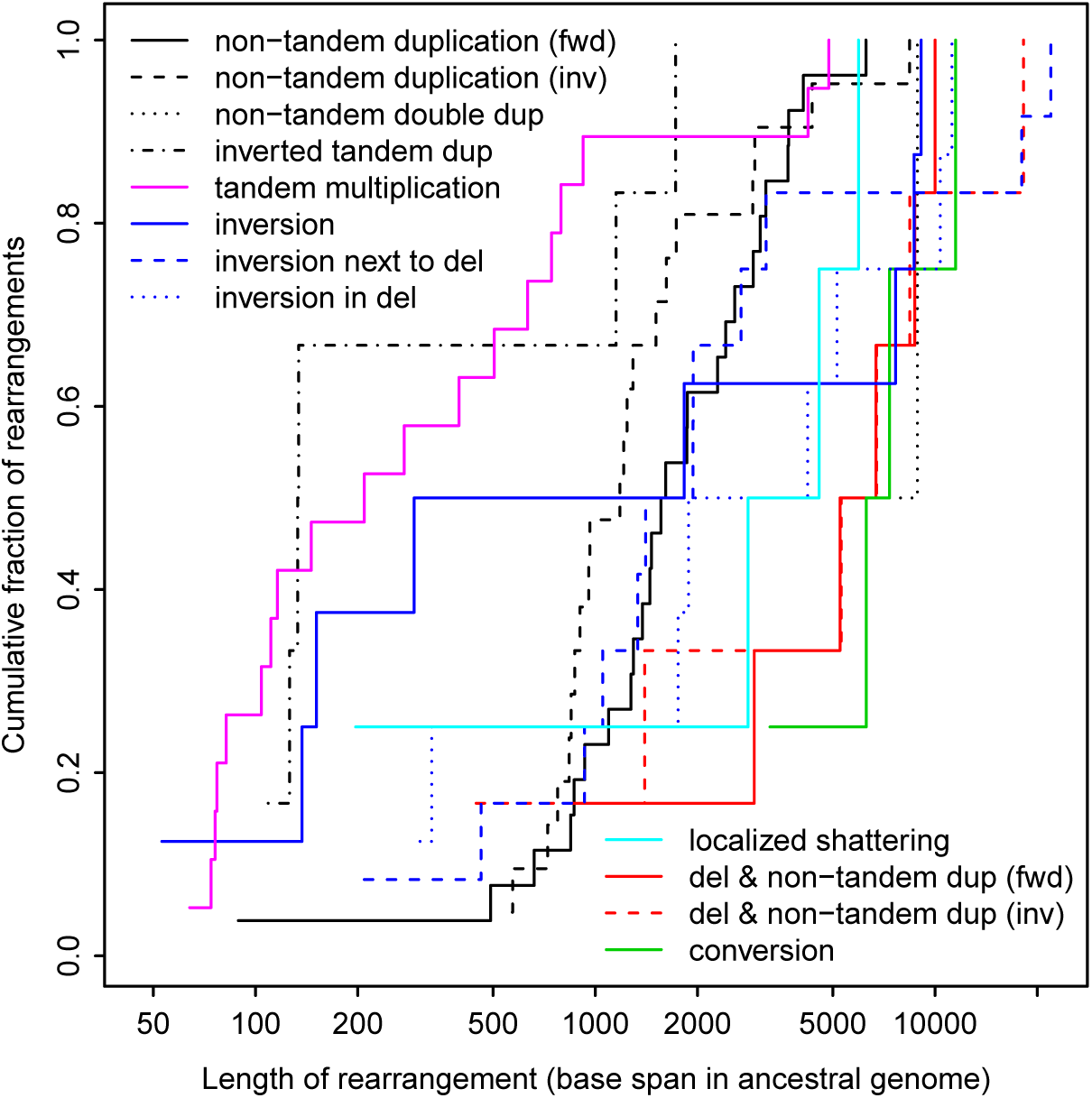
Lengths of rearrangements in the R9.4 dataset.

### Comparison to previously-found inversions

The P5-C3 reads are from a previous study [15], which published a table of 40 inversions. Fourteen of them match our rearrangements: 2× inversion, 6× inversion next to deletion, 3× inversion in deletion, 2× unique (Fig. 6 D–E), and 1× “dubious” (reference human genome not ancestral). So only a minority of each inversion catalog is common to both.

Seventeen of our inversions were not found in the previous study. The reason is unclear; however, among the 17 are our six longest (*>*8 kb) and three shortest (*<*200 bp) inversions.

We miss 26 of the previous inversions, for clear reasons. In 23 cases, the reference human genome lacks an unbroken alignment to chimp or gorilla, thus failing our ancestrality criterion. One other case could not be transferred between genome versions. (The previous study used an older reference human genome.) The remaining two cases are not “local” rearrangements. One of them (in chromosome 1q25) is basically an inversion, but a fragment of the DNA read next to the inversion aligns to a distal location. The other (in chromosome 9q34) does not look like an inversion: it seems to be an interchromosomal rearrangement involving chromosome 22.

Since we miss most of the previously-found inversions due to our ancestrality requirement, the reader may wonder whether this requirement is too strict. For inversions without deletion (Fig. 4I), it is indeed too strict: the alignments reliably indicate such inversions regardless of whether the reference genome’s arrangement is ancestral. For inversions with deletion (Fig. 4 J–N), however, ancestrality is important. Suppose the reference genome has an inversion with deletion, relative to the DNA reads. The regions of DNA reads that have been deleted in the genome should be unaligned, but if any part of the deleted sequence has a sufficiently-similar paralog anywhere else in the genome, the reads will be wrongly aligned to this paralog. Thus, while the ancestrality requirement could be relaxed for some types of rearrangement, in general it is necessary.

### Comparison to the Database of Genomic Variants

Finally, we compared the R9.4 rearrangements to structural variants in the Database of Genomic Variants (DGV release 2016-08-31) [34]. The boundaries of both our rearranged regions and DGV variants are not necessarily precise, so we sought overlaps. About 100 R9.4 rearranged regions (65%) roughly coincide with DGV variants (Fig. 8). By this criterion, DGV lacks most of our tandem multiplications (30/42) and inversions (6/8).

**Figure 8.**
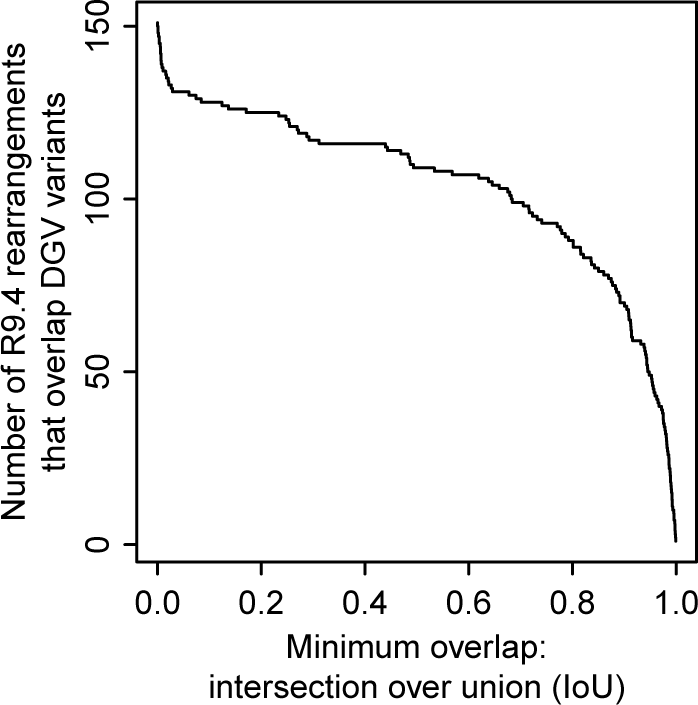
Number of R9.4 rearrangements that overlap variants in DGV.

Many R9.4 rearranged regions coincide near-exactly with DGV variants (Fig. 8), but the descriptions often differ. For instance, the double duplication in Fig. 4F coincides (intersection over union = 0.986) with variant esv3324, a “CNV loss”. This variant was found from short DNA reads [35]: it looks as though they correctly found the adjacency of the two duplicated fragments, but incorrectly interpreted this as deletion of the interval between the two duplication sources. This illustrates the advantage of longer reads, which reveal the whole rearrangement.

Diverse R9.4 rearrangements coincide with DGV “inversions”, including Fig. 4D (IoU 0.999), Fig. 4E (IoU 0.996), Fig. 4L (IoU 0.999), and Fig. 6F (IoU 0.99). These do all involve inverted fragments, but to describe e.g. Fig. 6F as “inversion” is at best incomplete and misleading. Short reads can show that neighboring parts of a sample genome map to opposite strands of a reference genome, but this is a feature of diverse types of rearrangement.

Further examples include Fig. 4M, which coincides (IoU 0.995) with a “CNV loss”, a correct but incomplete description. The conversions in Fig. 4S (IoU 0.938) and Fig. 4T (IoU 0.956) coincide with “CNV deletions”. Fig. 4B (IoU 0.981) and Fig. 6B (IoU 0.9) coincide with “tandem duplications”. The localized shattering in Fig. 4Q coincides (IoU 0.943) with a “CNV gain+loss”, which is consistent but vague.

In summary, our long read analysis clarifies previously-reported variants.

## Discussion

### Long DNA reads have many dubious rearrangements

Both datasets (R9.4 and P5-C3) have many orphan rearrangements, that occur in just one read and are not supported by other reads. A plausible explanation is that they are artifacts of the sequencing process. For R9.4, we can distinguish plausible from dubious rearrangements provided we have enough sequencing coverage that real rearrangements are expected to be represented in more than one read. For P5-C3, it was harder to distinguish plausible from dubious rearrangements, due especially to the dubious fountain-like rearrangements, which are large and often overlap other rearrangements.

Some, but not all, orphan rearrangements have characteristic patterns (e.g. zigzag, fountain-like) not seen in the supported rearrangements. So it may be possible to automatically recognize and discard them. This could be a moving target, however, if artifact patterns change with different versions of sequencing technology.

Another approach would be to automatically recognize if more than one DNA read has the same rearrangement, not merely overlapping rearrangements. This is not trivial, because two reads with the same rearrangement can have non-identical alignments due to sequencing error (e.g. Fig. 3C).

### Rearrangement patterns

Nearly all the supported rearrangements belong to recurring patterns/motifs. One pattern can manifest at a wide range of scales, e.g. hundreds of bp to tens of kb (Fig. 4 J–L).

The most common type of local rearrangement is tandem multiplication (especially duplication). This is very common in tandem repeats, as expected. Non-tandem duplications are the second-most frequent, but more complex patterns occur too, e.g. double duplication (Fig. 4 E–F), localized shattering (Fig. 4 Q–R), and a fascinating unique rearrangement (Fig. 6F). We have established here that such complex rearrangements can be detected in long DNA reads.

Gene conversions were also identified: although this is a classically-known phenomenon, it has surely been under-reported in high-throughput sequencing studies, due to the ambiguity in aligning these sequences. Probability-based alignment is a powerful way to resolve this kind of ambiguity.

Some patterns that might be expected do not occur. In particular, there are no cases of translocation (movement of a DNA fragment), without deletion or duplication.

None of the supported rearrangements have large unaligned “insertions” in the DNA reads (apart from cases like Fig 3C, which are likely alignment failures rather than true insertions). This fits the hypothesis that most sequence is descended from ancestral sequence, not spontaneously generated; it also supports the reality of these rearrangements. Thus the term “insertion” is potentially misleading: it is not symmetric with “deletion”, and is usually duplication (or deletion in the reference). On the other hand, orphan rearrangements often have unaligned parts of the reads (Fig. 2).

### Possible rearrangement mechanisms

A classic rearrangement-causing mechanism is homologous recombination. Some of the rearrangements are typical examples: conversion (Fig. 4 S–T) and inversion between oppositely-oriented repeats (Fig. 4H). Other rearrangements are harder to explain. The inversion in Fig. 4J lies between oppositely-oriented AluSx elements, suggesting recombination, but it is unclear how the adjacent deletion arose. The inversion in Fig. 6E overlaps oppositely-oriented Alus, but perhaps that is a coincidence: this inversion may be better explained by template switching during DNA replication.

It has been suggested that template switching by DNA polymerase, during DNA replication, is a cause of complex rearrangements. Template switching may be triggered by a broken replication fork, in a mechanism termed MMBIR (microhomology-mediated break-induced replication) [1], or a stalled replication fork, in a mechanism termed FoSTeS (fork stalling and template switching) [36]. The suggestion is that multiple template switches can occur, producing rearrangements with any number of duplications, deletions, and repositionings [1].

Some of the rearrangements are best explained by shattering of DNA into fragments, which then rejoined in random order and orientation, with some fragments lost (e.g. Fig. 4 Q–R). The main reason to favor this explanation over template switching is that some of the rearranged fragments come from adjacent parts of the ancestral sequence: we know of no reason why template switching would exhibit this tendency. Localized shattering and rejoining can be caused by radiation, such as cosmic rays and alpha particles from radon gas [37, 38, 39, 40].

The rearrangements are often associated with repeats. In some cases, this is expected: tandem repeats have high rates of duplication and conversion, similar repeats are prone to recombination, and simple repeats can be fragile. It is less clear, however, why rearrangement edges are often close to the edges of transposon fragments. The edge of a transposon fragment is necessarily the edge of a previous rearrangement or deletion which created that fragment: so maybe these sites are fragile.

### Aligning rearranged sequences

Although the Repeated Matches Algorithm was described decades ago (and `last-split` has been publicly available since 2013), its suitability for elucidating rearrangements has not been recognized. This algorithm simultaneously optimizes the division of a query sequence into (any number of) parts, and the alignment of each part to reference sequences, based on probabilities of substitution, insertion, and deletion. Here, we have demonstrated that it can find intricate rearrangements in long DNA reads, and handle ambiguous situations like gene conversion.

We have emphasized the importance of an ancestral reference sequence. The main problems with a non-ancestral reference are: reference-specific deletions and duplications. If there is a reference-specific deletion, the corresponding region of the query sequence should not be aligned, but if any part of it has a sufficiently similar paralog elsewhere in the reference, it will be aligned. Here, “sufficiently” similar is determined by the probabilities of substitution, insertion, and deletion used for alignment (Table 1). Thus, lower sequencing error rates would reduce this problem.

Our approach cannot align fragments shorter than some minimum length (which depends on the genome size and sequencing error rates). For example, it will not align a 12 bp fragment to a human genome, because such an alignment is too insignificant to distinguish from chance matches. However, it is possible to detect inversions *<* 12 bp [30]. The Repeated Matches Algorithm cannot do this because it assumes that rearranged fragments can be anywhere in the genome with uniform probability. Perhaps it could be modified to favor aligning nearby parts of a query sequence to nearby parts of the reference [41].

`last-split` is a useful approximation to the Repeated Matches Algorithm, but better approximations (in terms of accuracy and speed) are surely possible. One idea is to run the full Repeated Matches Algorithm on complex rearrangements tentatively identified by `last-split`.

### Ancestral genomes are ideal reference genomes

If we could reconstruct the most recent common ancestor of all extant human genomes, it would be an ideal reference for human nucleotide sequences. Almost every part of any human DNA read would be descended from a unique part of the reference. This would solve the worst problems with currently-used reference genomes.

### Identifying rearrangements from alignments

Although the alignment procedure can handle arbitrary rearrangements, this study only examined “local” rearrangements. A local rearrangement has a tightly-defined location: a start and an end coordinate in one chromosome. This property was used to: gather overlapping rearrangements in different DNA reads, check these regions’ alignments to chimp and gorilla, draw pictures of these regions, and compare them to DGV.

If we consider non-local rearrangements, even the concept of a discrete rearrangement becomes unclear. Rearrangement *events* are discrete, but they may interact to produce a complex pattern of sequence change. For example, two overlapping inversions produce a combined rearrangement that does not resemble two distinct rearrangements. We would ideally like to infer the history of rearrangement events from the final sequence, but this is a deep problem, especially if one event can be complex (e.g. shattering).

Our analysis pipeline is semi-manual: it makes pictures of rearranged regions for human interpretation. This is appropriate for exploring what kinds of (real and artifactual) rearrangements exist.

It would be convenient to automatically infer one or two rearranged alleles in each region. This is not trivial, because different DNA reads that cover the same rearrangement can have different alignments (e.g. Fig. 3C). This problem would be reduced with lower rates of sequencing error, because the different DNA reads would align more consistently (and presumably more accurately).

An alternative approach is to first perform reference-free assembly of the reads, then seek rearrangements in the assembly relative to a reference genome. It is not obvious which approach is better. Comparing unassembled reads to the reference genome is more direct: there is no question of assembly error, and we can examine the alignments of the original reads.

### Conclusions

This study demonstrates an approach to find arbitrarily-complex (but localized) sequence rearrangements in long DNA reads. Its key ideas are to exploit the asymmetric relationship between ancestral and derived sequences, and the use of probabilities to resolve ambiguities due to repeats and paralogs. The long reads studied here have many dubious (likely-artifactual) rearrangements, but it is possible to discriminate them from likely-real rearrangements. Almost all the real rearrangements belong to recurring motifs. Although simpler rearrangements are more common, complex and intricate rearrangements can be found: including localized shattering, which resembles radiation damage. Such rearrangements may be powerful homoplasy-free phylogenetic markers. It will be fascinating to survey many more genomes, from healthy and diseased cells, in a variety of organisms.

## Acknowledgements

We thank Risa Kawaguchi for suggestions on displaying rearrangements, and Anish Shrestha for comments on the manuscript.

